# Genomic and epigenomic maps of mouse centromeres and pericentromeres

**DOI:** 10.1101/2024.10.09.617447

**Authors:** Gitika Chaudhry, Jinguye Chen, Lucy Snipes, Smriti Bahl, Jenika Packiaraj, Xuan Lin, Jitendra Thakur

## Abstract

Satellite DNA makes up ∼11% of the mouse genome, predominantly located in centromeric and pericentric regions, which are crucial for chromosome segregation. While comprehensive assemblies of these regions have been established in the human genome, they are still lacking in the mouse genome. In this study, we used PacBio long-read sequencing, CUT&RUN sequencing, DNA methylation analysis, and RNA sequencing to generate genomic and epigenomic maps of these regions. We find that centromeric regions are primarily occupied by 120-mer Minor satellites, with other Minor Satellite length variants, 112-mers and 112-64-dimers, localized at centromere-pericentric junctions. Pericentromeric regions are mainly composed of homogeneous Major satellites, while pericentric-chromosomal junctions contain a higher density of divergent satellites. Additionally, the density of non-satellite repeats increases progressively from centromeres to pericentromeres, and further toward chromosomal arm junctions. We found that 120-mer Minor satellites in the core centromere are highly enriched with CENP-A, while the 112-mers and 112-64-dimers show lower CENP-A levels. Homogeneous Major satellites are more enriched with H3K9me3 heterochromatin, whereas divergent Major satellites are preferentially associated with H3K27me3. Furthermore, DNA methylation levels are lower in centromeres compared to pericentric regions. We also observed that only a small subset of satellites is transcribed into RNA, particularly regions exhibiting lower DNA methylation density. Our comprehensive assembly and characterization of the genomic and epigenomic landscape of mouse centromeric and pericentric regions have major implications for satellite biology and ongoing mouse telomere-to-telomere (T2T) assembly efforts.

## Introduction

Repetitive sequences are a fundamental component of eukaryotic genomes, comprising approximately 50% of the mammalian genome (de Koning et al., 2011). These sequences are crucial for maintaining the structural and functional integrity of the genome, characterized by their high copy number, and include various forms such as transposons, tandem repeats, and satellite DNA (Biscotti et al., 2015; Feschotte & Pritham, 2007; Thakur et al., 2021). Satellite DNA consists of tandemly repeated sequences often located in specific chromosomal regions, such as centromeres, pericentromeres, and telomeres, where they play essential roles in maintaining chromosomal stability, facilitating chromosome segregation during cell division, and protecting chromosome ends (Alexandrov et al., 2001; Biscotti et al., 2015; Garrido-Ramos, 2017; Plohl et al., 2012; Rudd et al., 2006; Thakur et al., 2021). Satellite DNA constitutes ∼11% of the mouse genome and ∼6-8% of the human genome (Altemose et al., 2022; Komissarov et al., 2011; Thakur et al., 2021). Centromeric satellites are present at the primary constriction, where they assemble specialized nucleosomes in which canonical histone H3 is replaced by its variant called centromeric protein A (CENP-A) (Ando et al., 2002; Henikoff et al., 2015). The CENP-A chromatin lays the foundation for the formation of the kinetochore, a protein complex critical for the attachment of chromosomes to spindle microtubules during mitosis and meiosis (Aldrup-Macdonald & Sullivan, 2014; Alexandrov et al., 2001; Altemose et al., 2022). Pericentromeric regions, which flank the centromeres, are also rich in satellite sequences and are bound to cohesin, which is protected from degradation until anaphase onset to ensure faithful chromosome segregation (Altemose, 2022; Guenatri et al., 2004; Hahn et al., 2013). Despite their functional importance, the assembly of these satellite regions has been challenging due to their highly repetitive nature. Recent advances in long-read sequencing technologies have led to significant improvements in genome assemblies, resulting in a complete end-to-end assembly for the human genome.

In humans, centromeres are composed of α-satellite DNA, which consists of tandem repeats of 171 base pairs forming higher-order repeat structures crucial for centromere function (Aldrup-Macdonald & Sullivan, 2014; Alexandrov et al., 2001; Rudd & Willard, 2004). These α-Satellite DNA can either exist as monomers, which are inactive and pushed toward pericentric regions or as higher-order repeats, in which a set number of monomers (ranging from 2 to 34) are repeated several times and can occur as active or inactive regions of centromeres (Alexandrov et al., 2001; Hartley & O’Neill, 2019; Waye et al., 1987). Human pericentric regions contain various satellite families (e.g., HSATI, HSATII, HSATIII, β-satellites), each distinct in their DNA sequences (Greig & Willard, 1992; Waye & Willard, 1989) (Altemose, 2022; Thakur et al., 2021). Unlike humans, where centromeres exhibit high variation in α-satellites and pericentric regions contain multiple satellite families, mouse centromeres are composed exclusively of Minor Satellites (MiSats), characterized 120 base pair repeat units, and pericentric regions consist solely of Major Satellites (MaSats), characterized by 234 base pair repeat units (Packiaraj & Thakur, 2024; Thakur et al., 2021). Human centromeric and pericentric regions exhibit high global and local variations, with repeat units in a given satellite class sharing approximately 60-100% sequence similarity (Aldrup-Macdonald & Sullivan, 2014; Altemose et al., 2022; Nurk et al., 2022). In addition, the human genome contains several chromosome-specific α-satellite sequences and higher-order repeat structures (Altemose et al., 2022; Nurk et al., 2022). These sequence variations have allowed the Telomere-to-Telomere (T2T) consortium to assemble a complete, gapless sequence, providing a comprehensive map of centromeric, pericentric, and telomeric regions (Altemose et al., 2022; Nurk et al., 2022). In contrast, mouse centromeric and pericentric satellites were previously considered highly homogeneous, and assembly maps have not been attempted. Thus, despite being a widely used model organism for genetic and biomedical research, the mouse genome still lacks a complete assembly map for these repetitive regions. Recently, we characterized the local arrangement of centromeric and pericentric satellites and the enrichment pattern of associated chromatin on PacBio HiFi reads (∼15 kb long) (Packiaraj & Thakur, 2024). We demonstrated the presence of considerable sequence variations within both centromeres and pericentric satellite regions in mouse (Packiaraj & Thakur, 2024). Although the extent of these variations is much lesser than human satellites, they can potentially contribute to the assembly of more extended maps for mouse centromere and pericentric regions.

In this study, we created detailed representative genomic and epigenomic maps (up to ∼1.4 Mb long) for the centromeric, pericentric, and junction regions in mouse reference C57BL/6J strain. We generated long-read sequencing reads on the PacBio Revio system and assembled the resulting HiFi sequencing reads into long contigs using the Hifiasm assembler. We have developed comprehensive and extensively annotated maps that encompass centromeres and pericentromeres, as well as telomere-centromere junctions, centromere-pericentromere junctions, and pericentromere-chromosomal arm junctions (extending up to approximately 1.5 Mb in length). Additionally, we have generated high-resolution chromatin profiling data for CENP-A, sequence-specific centromeric protein CENP-B, H3K9me3, and H3K27me3 chromatin to understand the enrichment patterns on these satellite assembly maps. We also analyzed DNA methylation profiles using kinetics tags of PacBio reads. Furthermore, we generated and analyzed RNA sequencing data from C57BL/6J mouse tissues to understand the transcriptional readouts from these regions. These findings provide critical insights into the complex dynamics of satellite DNA and its impact on genomic function and stability and contribute substantially to ongoing mouse T2T assembly efforts.

## Results

### Assembly maps of mouse centromeres and pericentric regions

To investigate the arrangement patterns of satellites on centromeres and pericentromeres on large assemblies, we generated HiFi long-read sequencing data utilizing the PacBio Revio sequencing platform from a male C57BL/6J mouse kidney tissue. The PacBio reads were assembled using the Hifiasm assembler (Cheng et al., 2021), and the resulting contigs were annotated using RepeatMasker with the *Mus musculus* repeat database and NCBI BLAST, employing satellite consensus sequences as databases. We isolated and analyzed contigs spanning centromeric MiSats, pericentric MaSats, centromeric-telomeric junctions, and pericentric-chromosomal arm junctions.

#### Centromeric maps

We identified 14 contigs with >5 kb long stretches of continuous MiSat arrays for centromeric regions, ranging from 63 kb to 1.16 Mb (Figure 1A & 1C). Seven centromeric contigs contained telomeric and pericentric junctions on opposite ends, and the distance from telomeric to MaSat-containing ends varied from 690 kb to 850 kb (average length=745 kb, n=7). We identified a 90 kb centromeric sequence on a contig with genes known to be present on the centromeric end of Y-chromosomes (Supplementary Figure 1). This 90 kb Y-centromere sequence is similar to the mouse Y-centromere previously mapped to a 90 kb region using a BAC clone (Pertile et al., 2009). Interestingly, the Y-centromere lacked flanking pericentric MaSats on the ∼1.2 Mb contig (Figure 1C). The ∼166 kb BAC clone in which the Y-centromere was initially identified also lacked flanking MaSats (Pertile et al., 2009). Such a lack of MaSats near Y-centromeres was also observed during the cytological visualization of mouse male chromosome spreads probed with MaSats (Falconer et al., 2010; Pertile et al., 2009). We also created Hifiasm assemblies from publicly available long-read sequencing data from a C57BL/6J female (Hon et al., 2020), which contained three contigs with both telomeric and pericentric ends with centromere lengths ranging from 717 kb to ∼1Mb (Supplementary Figure 1). These findings indicate that mouse centromere size ranges from 90 kb to ∼1 Mb (Supplementary Figure 1).

**Figure 1.**
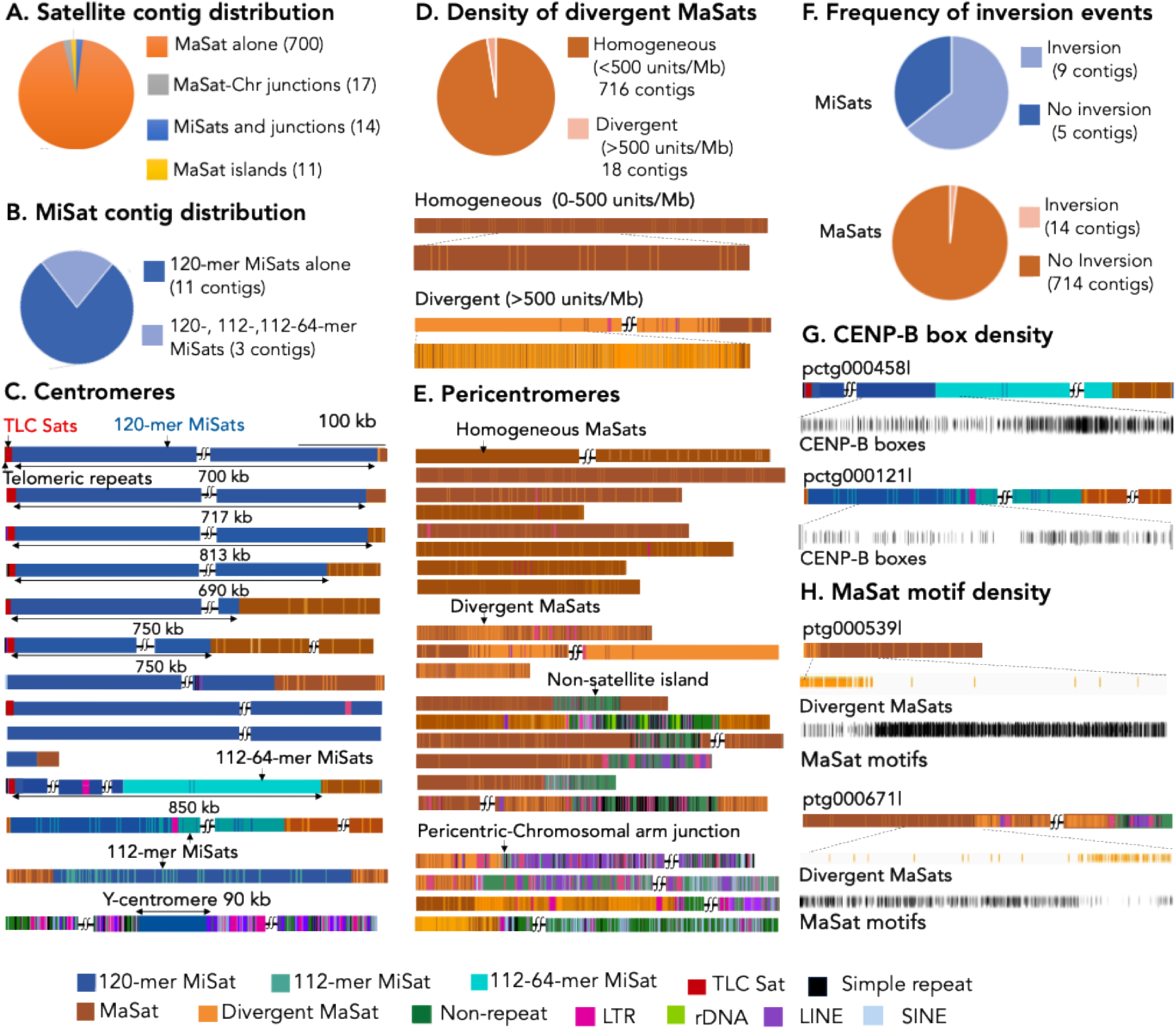
Arrangement of centromeric and pericentric satellites on assembled HiFi contigs. **A)** Number and distribution of different satellite containing contigs **B)** Number and distribution of centromeric contigs based on the type of MiSat length variants. **C)** All contigs containing > 5 kb long stretches of MiSat units. **D)** Number and distribution of pericentric contigs based on the density of divergent MaSats (Top). Zoomed in view of pericentric regions based on the density of divergent MiSats (Bottom). **E)** Representative contigs from pericentromeres and pericentric-chromosomal arm junctions. Contigs are drawn to scale, and a size bar is given. All contigs for centromeric and pericentric regions are shown in Supplementary Figures 1 and 2, respectively. **F)** The distribution of contigs based on the presence and absence of inversion events. **G)** CENP-B box density on centromeric contigs. A 150 kb region is zoomed in from the indicated regions for each contig. **H)** MaSat motif box density on pericentric contigs. A 150 kb region is zoomed in from the indicated regions for each contig.

As expected, centromere-telomere junctions were occupied by L1 LINE elements and TLC satellites (Figure 1C). In addition to 120-mer MiSats, mouse centromeres contain 112-mer and 112-64-dimer MiSat length variants (Packiaraj & Thakur, 2024). We found that 120-mer MiSats were present on all MiSat contig types, but 112-mer and 112-64-dimer MiSats were only detected in 2 and 1 contigs, respectively (Figure 1B &1C). Interestingly, 112-mer and 112-64-dimer MiSats were always present at regions adjacent to MaSats, suggesting that these MiSat length variants are specific to centromere-pericentric junctions of certain chromosomes (Figure 1C, Supplementary Figure 1). Both contigs containing 112-mer MiSats also contained MaSats on both sides, indicating possible rearrangements leading to the insertion of pericentric sequences at these MiSat variants near centromere ends (Figure 1C & Supplementary Figure 1), similar to centromeres of human chromosomes 3 and 4. We also created Hifiasm assemblies from publicly available long-read sequencing data from a C57BL/6J female (Hon et al., 2020). Similar to our findings in the male, we detected two 112-mer and one 112-64-dimer MiSat contigs in the female dataset (Supplementary Figure 1), ruling out the possibility that these MiSat variants are specific to Y-chromosome centromeric junctions.

#### Pericentric maps

We identified 734 contigs for pericentric regions consisting of long continuous arrays of MaSats ranging up to ∼1.4Mb in length (Figure 1D, Supplementary Figure 2). This large number of contigs is expected because MaSats make up about 10% of the mouse genome, compared to approximately 1% for MiSats. We have previously shown that ∼10% of MaSat containing HiFi long reads from the C57BL/6J strain contain divergent MaSat units that share <75% sequence similarity to the MaSat consensus sequence (Packiaraj & Thakur, 2024). Here, we found that while the majority of contigs contained predominantly homogeneous MaSats, with divergent MaSats interspersed at low density (<500 satellite units/Mb), a small fraction (∼3.2%) of MaSat contigs contained divergent MaSats at a high density (> 500 satellite units/Mb) (Figure 1D, Supplementary Figure 2). All 12 pericentric-centromeric junction contigs that were also identified as MiSat-containing contigs contained homogeneous MaSats (Figure 1C, Supplementary Figure 1).

We determined the distribution of known sequence motifs in centromeric and pericentric regions. The specific sequence motif found in MiSats is the CENP-B boxes, while MaSats contain the Major motif (Ohzeki et al., 2002; Packiaraj & Thakur, 2024). The CENP-B boxes are essential for recruiting and binding to the centromeric protein CENP-B, but the function of the Major motif is still not understood. We found local variations in CENP-B box density in MiSat contigs (Figure 1G). Interestingly, the density of CENP-B boxes was highest on 112-64 dimeric MiSats and lowest in 120-mer MaSats interspersed with low-density 112-mers. In 112-mer containing contigs, the density of 112-mers and CENP-B boxes increased progressively toward centromeric-pericentric junctions (Figure 1G). The Y-centromere lacked CENP-B boxes (Not Shown). In pericentric regions, the density of the Major motif was inversely proportional to the density of divergent MaSats. The highest density was found in homogeneous MaSats, while the lowest density was observed in high-density divergent MaSats (Figure 1H).

We identified inversion events in satellite contigs by analyzing the directional changes in the CENP-B motif in MiSats and the MaSat motif in Major Satellites. Inversions were highly frequent in centromeric regions, with 9 out of 14 centromeric contigs exhibiting these inversions, suggesting frequent structural rearrangements within these regions (Supplementary Figure 4A and 4B). In contrast, inversion events were less frequent in pericentric regions, appearing in only 14 out of 700 contigs. In homogeneous MaSat contigs, inversions were primarily located near interspersed non-satellite sequences, suggesting a possible association between inversion events and the presence of non-satellite DNA elements (Supplementary Figures 4A and 4B).

Overall, the frequent occurrence of inversion events in centromeric regions suggests that they are more prone to chromosomal rearrangements than adjacent pericentric regions.

### Divergent MaSats are preferentially localized at pericentric-chromosomal arm junctions

Out of the 734 MaSat-containing contigs analyzed, we identified 19 in the pericentric-chromosomal junction regions, each containing several Mb long chromosomal arm sequences (Figure 2A, Supplementary Figure 3). To determine the chromosomal origin of these contigs, we mapped annotated genes located near chromosomal ends in the mm39 genome assembly to these contigs. This allowed us to assign 17 of the 19 pericentric-chromosomal junction contigs to specific chromosomes (Figure 2A). As discussed in previous sections, the contig associated with the Y-centromere did not contain any MaSat sequences.

**Figure 2:**
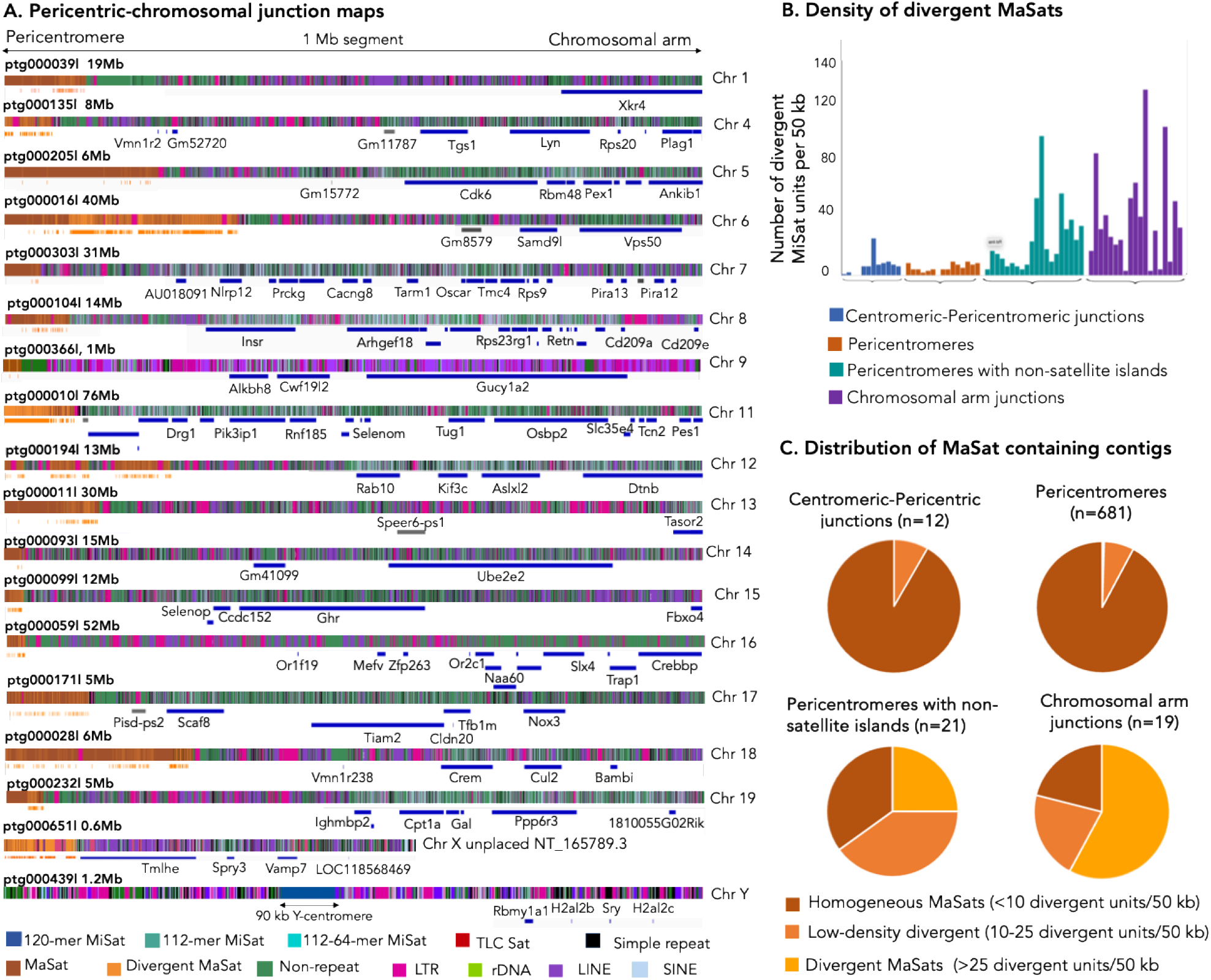
Pericentric-chromosomal junctions are enriched for divergent MaSats. **A)** Mapping of pericentric-chromosomal arm junctions. For each contig containing a pericentric-chromosomal junction, a 1 Mb region is displayed, showing both the chromosome-specific genes that map to the mm39 genome assembly and the junction contig itself. Not all genes shown in the figure are named due to space constraints. A complete list of these genes is given in Supplementary Figure 3. Below each contig map, the divergent MaSat units are marked in orange, annotated genes are represented in blue, and pseudogenes are shown in grey. **B)** Divergent MaSat density across various MaSat-containing regions. Density of MaSats in contigs from centromeric-pericentric regions, pericentric regions alone, pericentric regions containing non-satellite islands, and pericentric-chromosomal arm junctions. For pericentric alone regions, 15 random contigs are included in the analysis. **C)** Distribution of indicated MaSat containing contigs based on the density of MaSats.

Among the 20 pericentric-chromosomal junction contigs, 16 contained high-density divergent MaSat sequences localized adjacent to the chromosomal arm regions. This preferential localization suggests that high-density divergent MaSats are primarily found near pericentric-chromosomal junctions (Figure 2, Supplementary Figure 2). Additionally, we identified 10 MaSat contigs interspersed with a ∼100-200 kb non-satellite island (Figure 1A, Supplementary Figure 2), as well as an additional 10 MaSat contigs that contained non-satellite sequence segments (<100 kb) at one end. Unlike pericentric-chromosomal junction contigs containing several Mb of non-satellite regions that could be assigned to specific chromosomes, the contigs with smaller (<100 kb) non-satellite regions could not be assigned to any chromosomal arm junctions, and these smaller non-satellite segments likely represent interspersed islands within the pericentric regions.

We calculated the density of divergent MaSat units in various categories of MaSat contigs, including those from centromeric-pericentric junctions, pericentric regions alone, pericentric regions containing non-satellite islands, and pericentric-chromosomal arm junctions. We found that the density of divergent MaSat units was highest in the pericentric-chromosomal junction contigs (Figure 2B-2C). These findings parallel those observed in human pericentromeres, where older, divergent sequences tend to localize toward the chromosomal arms, while recently evolved homogeneous sequences remain closer to the core centromeres (Altemose et al., 2022; Nurk et al., 2022).

### Retrotransposon density increases gradually from centromeres to chromosomal arms

We determined the density and arrangement of non-satellite repetitive sequences, including LTR retrotransposons, non-LTR retrotransposons such as LINEs (long-interspersed nuclear elements) and SINEs (short-interspersed nuclear elements), as well as simple repeats on centromeric and pericentric contigs. In the centromeric regions, we identified four contigs, each containing a single interspersed LTR retrotransposon belonging to the ERVK family (Figure 3A-B, Supplementary Figure 1). Specifically, three of these contigs contained IAP (Intracisternal A-particle), while one contained the ERVB7_1-LTR_MM element (Figure 3A-B, Supplementary Figure 1). Centromeric regions were largely devoid of SINEs, LINEs, and simple repeats (Figure 3A-3C). We only found one contig containing two simple repeats and another containing three SINE (B2_mm1t) elements (Figure 3A-3C). No LINE elements were detected within MiSat contigs. We confirmed the presence of telomeric repeats, categorized as simple repeat sequences at the telomeric end of centromeric contigs (Figure 3A). Additionally, we observed L1 LINE elements at the centromeric-telomeric junctions (Figure 3A), consistent with previous reports (Kalitsis et al., 2006).

**Figure 3:**
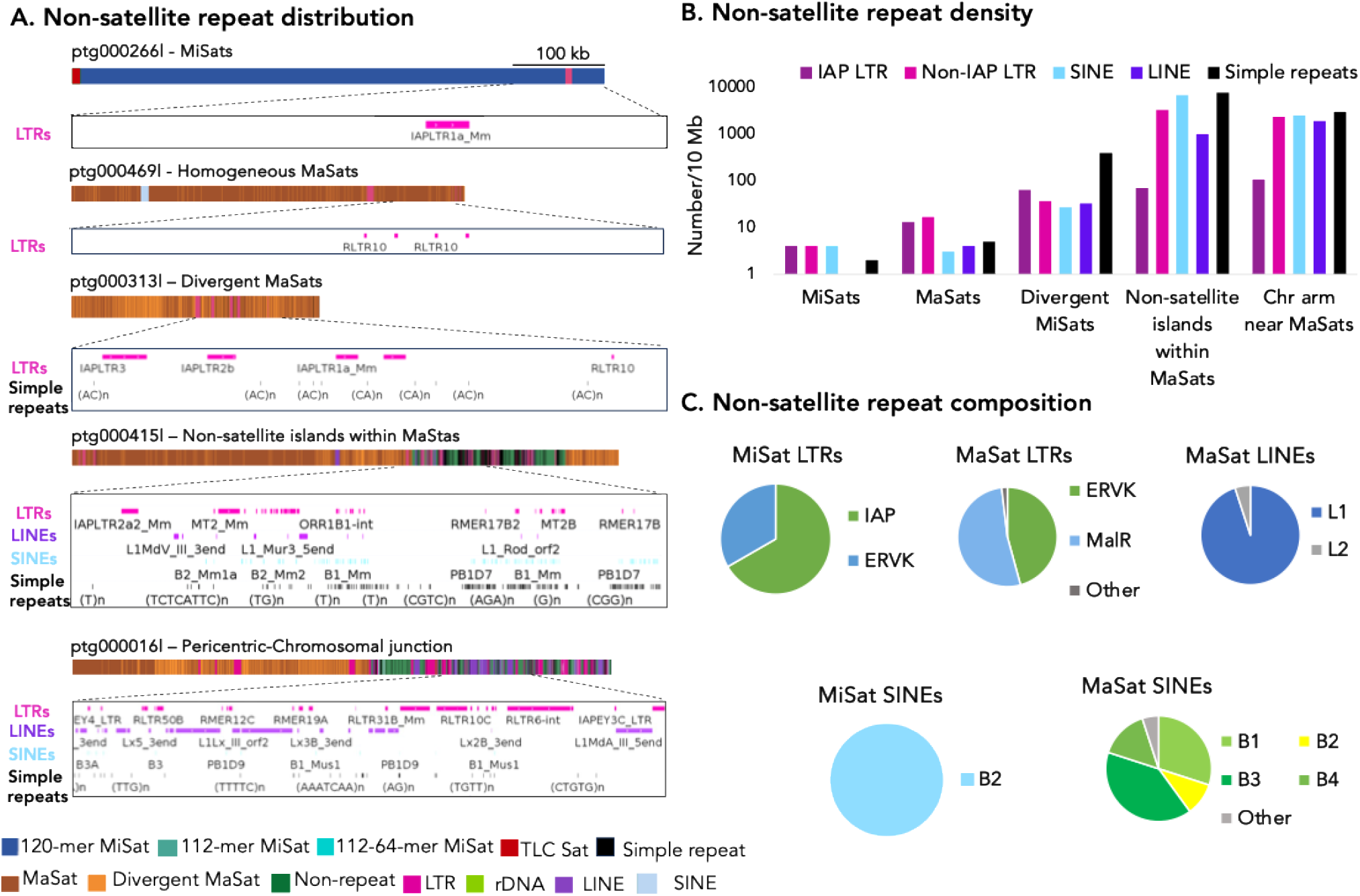
Non-satellite repeat composition at centromeric and pericentric regions. **A)** Representative contigs showing the distribution of LTRs, LINEs, SINEs, and simple repeats on centromeric, pericentric, and junction regions. Contigs are drawn to scale. A zoomed-in view across a 100 kb region is shown below each contig. **B)**. The density of retrotransposons (IAP and non-IAP LTRs), SINEs, LINEs, and simple repeats across satellite regions. **C)**The composition of SINEs, LINEs, and simple repeats on centromeric MiSats, pericentric MaSats, and Chromosome arm junctions.

The density of LTR retrotransposons, LINEs, SINEs, and simple repeats increased in pericentric MaSat contigs (Figure 3A-3C). In pericentric MaSats, we observed the presence of IAPs and other ERVK family members, such as RLTRs. These pericentric regions also contained MalR LTRs from the ERVL family (Figure 3A-B, Supplementary Figure 3). Unlike centromeric MiSats, which only contained B2 SINE elements, pericentric MaSats contained B1, B2, B3, B4, and others. (Figure 3A-3C). LINE elements in the pericentric MiSats were from the L1 and L2 categories (Figure 3C). All retrotransposons and simple repeats were more densely distributed on divergent MaSats than homogeneous MaSats, likely because divergent MaSats are predominantly located near chromosomal arm junctions.

MaSats in pericentric junction proximal to chromosomal arms contained a higher density of LTR retrotransposons, LINEs, SINEs, and simple repeats than centromeric and pericentric contigs (Figure 3A & 3B). Furthermore, the density of all non-satellite repeats was highest in non-satellite islands within MaSats (Figure 3A & 3B). This suggests that these pericentric-chromosomal arm junctions and islands are hotspots for non-satellite repeat integration. We found a few tRNA genes at pericentric major-chromosome transition zones (Figure 3A). Such elements act as barrier elements at the pericentric heterochromatin in fission yeast (Kuhn et al., 1991; Scott et al., 2006), and they may have a similar function in mouse pericentric regions.

Overall, the density of simple repeats, retrotransposons, LINEs, and SINEs increased gradually from centromeric regions to pericentric-chromosomal arm junctions. This progressive increase suggests a structured arrangement of non-satellite repeat sequences, with the highest density at the transition to chromosomal arms. This gradient of non-satellite repeat density potentially indicates regions of increased genomic rearrangements or structural transitions, underscoring the increasing complexity of repetitive sequences as one moves away from the centromere.

### Centromeric and pericentric regions and junctions show distinct chromatin enrichment patterns

To explore how the sequence composition and arrangement of satellite variants across different contigs influence chromatin organization, we assessed the enrichment of key chromatin marks and centromeric proteins in centromeric and pericentric contigs. We generated high-resolution CUT&RUN sequencing data from C57BL/6J mouse tissue for CENP-A, CENP-B, H3K9me3, H3K27me3, and an IgG control. Mapping the CENP-A CUT&RUN data to Hifiasm contigs revealed that contigs containing 120-mer MiSats exhibited the highest levels of CENP-A enrichment, while contigs containing 112-mers and 112-64-dimers showed lower enrichment (Figure 4A). This observation aligns with our previous observations of CENP-A binding patterns in HiFi reads (Packiaraj & Thakur, 2024). CENP-A levels were markedly reduced on TLC satellite sequences and absent on telomeric repeats (Supplementary Figure 5). Furthermore, IAP elements interspersed within MiSats showed no CENP-A enrichment (Supplementary Figure 5) as previously shown (Packiaraj & Thakur, 2024).

**Figure 4.**
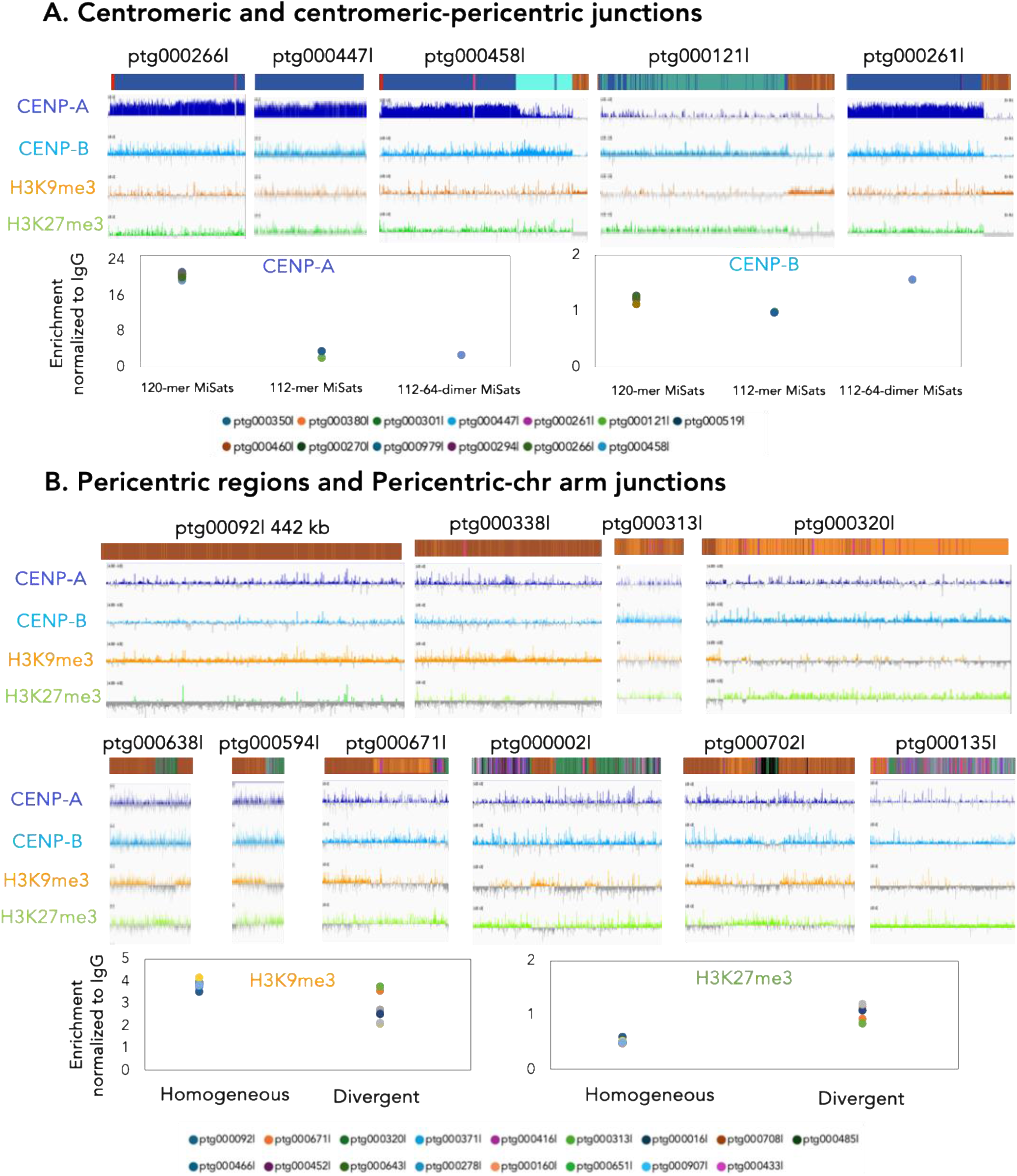
Enrichment of chromatin and centromeric proteins on centromeric and pericentric regions. **A)** Chromatin profiles showing the log2 enrichment ratio of CENP-A, CENP-B, H3K9me3, and H3K27me3 compared to an IgG control across centromeric and centromeric-pericentric junction regions (Top panels). Enrichment signals were quantified by normalizing the number of CENP-A and CENP-B reads mapped to 120-mer, 112-mer, 112-64-dimer, and MaSat-containing regions from centromeric contigs against the IgG control (Bottom panels). **B)** The log2 enrichment ratio of CENP-A, CENP-B, H3K9me3, and H3K27me3 is shown for centromeric and centromeric-pericentric regions (Top panels). Similarly, the number of H3K9me3 and H3K27me3 reads mapped to homogeneous and divergent MaSat-containing regions from pericentric contigs were normalized against the mapped IgG control reads (Bottom panels). The Y-axis range is consistent across all tracks (-4 to +4). Contigs are drawn to scale.

Interestingly, despite lower CENP-A enrichment, the 112-64-dimer MiSats showed the highest enrichment of CENP-B among all analyzed satellite contigs (Figure 4A). Similarly, although 112-mer-containing contigs had reduced CENP-A levels, their CENP-B enrichment was comparable to that of 120-mer-containing contigs (Figure 4A). The 112-mer and 112-64-dimer MiSats contain a higher density of CENP-B boxes (Figure 1G). Given that CENP-B binds in a sequence-dependent manner (Fachinetti et al., 2015), high levels of CENP-B enrichment, independent of CENP-A on these sequences are expected. Besides its well-known role in centromeric function, mouse CENP-B has been implicated in pericentric H3K9me3 heterochromatin formation (Kumon et al., 2021). However, the mechanism by which a centromere-specific protein like CENP-B contributes to pericentric heterochromatin remains unclear. Our observation of high CENP-B enrichment in centromeric-pericentric boundary regions suggests that CENP-B may interact with pericentric junction-specific proteins to play a role in H3K9me3 heterochromatin formation.

Next, we mapped the CENP-A, CENP-B, H3K9me3, H3K27me3, and IgG control CUT&RUN data to pericentric MaSat contigs. We found that homogeneous MaSats are consistently enriched with H3K9me3 across long continuous stretches of contigs, while high-density divergent MaSats show a more discontinuous enrichment of H3K9me3. Notably, the densest divergent MaSat regions exhibited the lowest H3K9me3 enrichment (Figure 4B). These divergent MaSats, however, were enriched with higher levels of H3K27me3 compared to homogeneous MaSats (Figure 4B). Similarly, non-satellite islands within MaSats displayed reduced H3K9me3 enrichment and increased H3K27me3 levels compared to the surrounding MaSats (Figure 4B). These findings suggest that high-density divergent MaSats and non-satellite islands are associated with reduced H3K9me3 heterochromatin and increased H3K27me3 enrichment, indicating distinct epigenetic regulation in these regions.

### Pericentric regions exhibit denser DNA methylation than centromeric regions

In humans, centromeric regions show a pronounced dip in DNA methylation compared to flanking pericentric regions. We assessed CpG DNA methylation in C57BL/6J satellite contigs by analyzing the kinetic signatures associated with methylated bases in PacBio HiFi reads. Both DNA methylation and CpG density were significantly higher in pericentric contigs than in centromeric regions (Figure 5A-5E). However, the reduction in DNA methylation in centromeric contigs compared to pericentric regions was less pronounced than what has been observed in humans (Altemose et al., 2022). Among the pericentric MaSats, homogeneous MaSats exhibited denser DNA methylation than divergent MaSats (Figure 5D). However, the methylation pattern was not entirely uniform, as pockets of lower DNA methylation, similar to those found in high-density divergent MaSats, were also present in homogeneous MaSats (Figure 5C).

**Figure 5.**
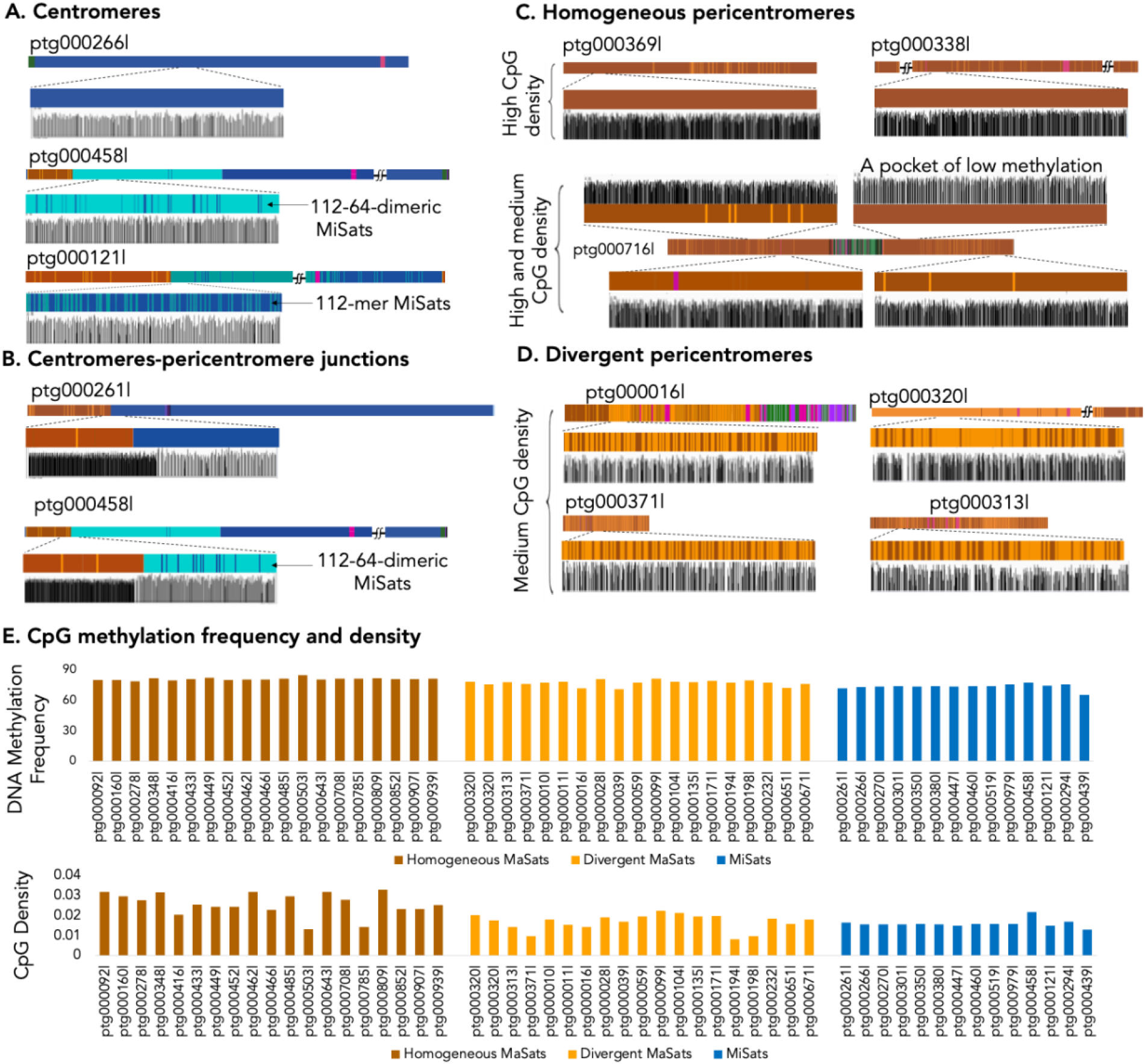
Pericentric regions show high levels of DNA methylation. CpG methylation frequency and density tracks on **A)** centromeric, **B)** centromeric-pericentric junction, **C)** homogeneous pericentric, and **D)** Divergent pericentric contigs are shown. A zoomed-in view across a 25 kb region is shown. Contigs are drawn to scale. **E)** Quantification of CpG DNA methylation frequency (Top) and CpG density (Bottom) on homogeneous MaSats, divergent MaSats, and MiSats containing contigs.

### Only a small subset of satellite repeats transcribes into RNA

Both centromeres and pericentromeres have been shown to transcribe in humans (McNulty et al., 2017; Velazquez Camacho et al., 2017). In contrast, studies in mice have predominantly focused on the pericentric regions, where non-coding RNAs transcribe from MaSats (Probst et al., 2010; Velazquez Camacho et al., 2017). To identify which centromeric and pericentric satellites are transcribed and to compare their expression levels, we performed RNA sequencing (RNA-seq) on total RNA extracted from C57BL/6J liver tissue and mapped the reads to satellite contigs. Our results revealed that only a small subset of MiSat and MaSat sequences were transcribed, and their expression levels were significantly lower—by several orders of magnitude—compared to the expression of rDNA genes interspersed within satellite contigs (Figure 6A-6C), suggesting that satellite non-coding regions are transcribed at very low levels. Notably, none of the contigs with a high density of divergent MaSats showed detectable RNA expression (Figure 6A). In regions of homogeneous MaSat contigs where transcription occurred, we observed less dense DNA methylation than the average methylation levels across homogeneous MaSat regions (Figure 6B). This indicates that MaSat transcripts likely arise from localized pockets of reduced DNA methylation within homogeneous pericentric regions. Furthermore, interspersed elements within both homogeneous MaSats exhibited higher expression levels than those within high-density divergent MaSats (Figure 6A and 6C). We did not observe a significant relationship between DNA methylation frequency and RNA expression levels (Figure 6D, Top panel). However, contigs with high RNA expression levels were associated with low CpG methylation density (Figure 6D, Bottom panel).

**Figure 6.**
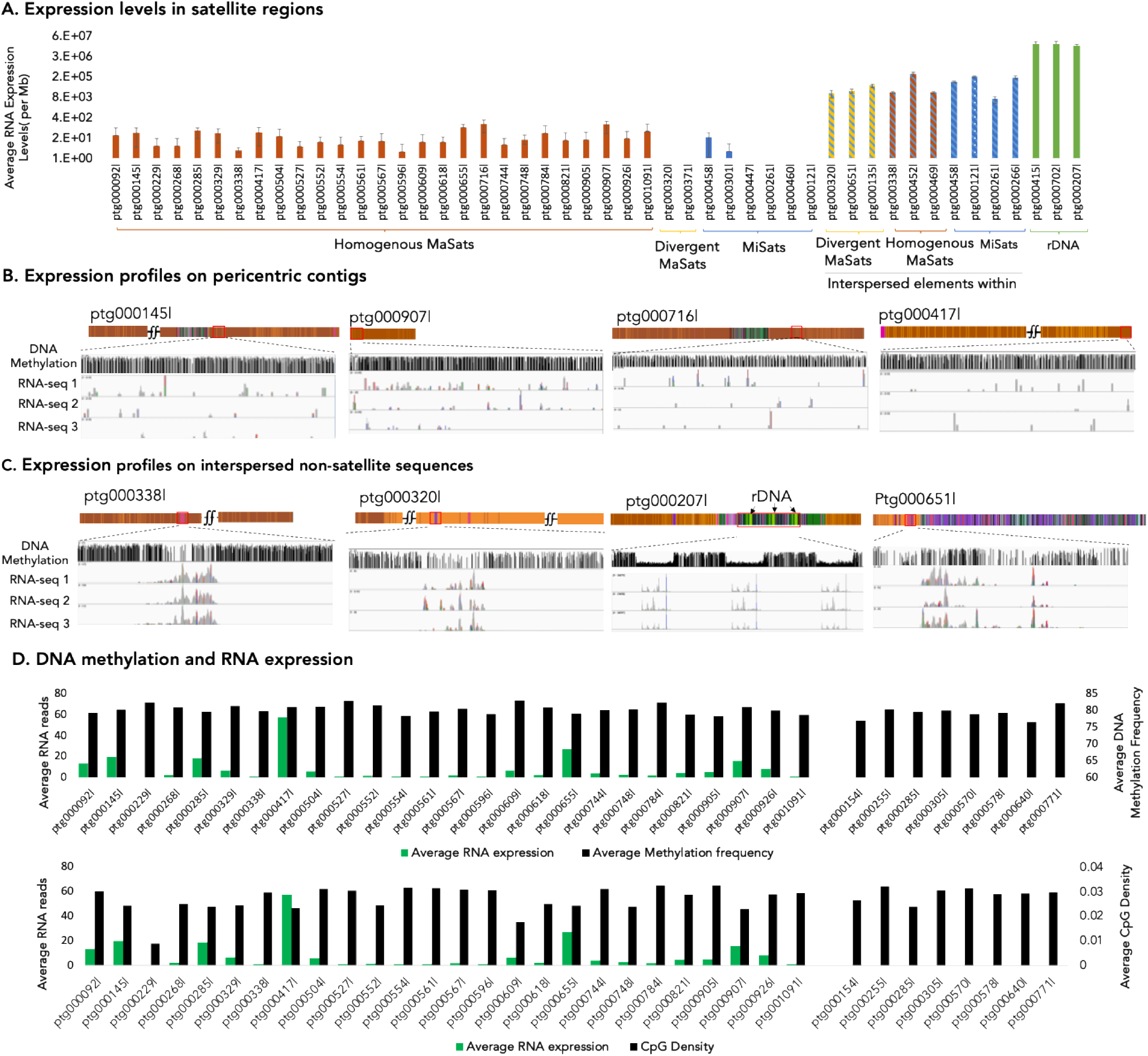
Transcription profiles of centromeric and pericentric regions. **A**) Expression levels on satellite contigs. The number of reads from three RNA-seq experiments from C57B6/J are mapped to indicated contigs and normalized to the length of the contig. RNA-seq profiles on **B)** pericentric contigs and **C)** non-satellite sequences interspersed within satellite contigs. Zoomed-in views across a 20 kb region from example contigs are shown. **D)** Average DNA methylation frequency (Top) and the CpG density (Bottom) on indicated homogeneous MaSat contigs are plotted along with the average RNA expression levels.

## Discussion

Satellite DNA constitutes approximately 11% of the mouse genome, predominantly located in the centromeric and pericentric regions, where it plays a crucial role in accurate chromosome segregation during cell division. Despite their critical importance, satellite DNA regions have historically posed challenges for genome assembly and annotation due to their highly repetitive nature. As a result, current mouse genome assemblies lack detailed maps of these regions, highlighting the need to fill the gaps in satellite DNA regions. In this study, we have created genomic and epigenomic maps of mouse centromeres and pericentric regions using long-read sequencing and epigenomic profiling assays.

Our findings suggest that in the reference laboratory C57BL/6J strain, centromeres can vary in length from 90 kilobases to ∼1Mb, similar to humans, where centromeres range from 500 kb to 5 Mb (Altemose et al., 2022). Most centromeric contigs contain 120-mer MiSats, while variants 112-mers and 112-64-dimer MiSats are predominantly found in regions adjacent to pericentric regions of a few chromosomes. These variants are present in male and female C57BL/6J animals, indicating that they are not specific to the Y-centromere. Interestingly, we confirmed that the Y-centromere lacks flanking pericentric MaSats, which agrees with previous findings suggesting that the majority of mouse Y chromosome is euchromatic (Morgan & Pardo-Manuel de Villena, 2017; Soh et al., 2014). Future studies investigating the loss rate of mouse Y-chromosome may shed light on whether the absence of pericentric MaSats reduces the chromosome segregation efficiency of Y-chromosomes.

Our results also show that most pericentric contigs, including those spanning centromeric-pericentric junctions, are populated mainly by homogeneous MaSats, while divergent MaSats preferentially localize toward the pericentric-chromosomal junctions, suggesting that, as in humans, sequence divergence of pericentric satellites increases with distance from the core centromere. Additionally, while centromeric regions contain rare short, interspersed transposons (a few kb in length), pericentric regions contain both short transposons and long islands of non-satellite sequences (ranging from ∼100 to ∼200 kb), indicating that pericentric regions are more permissive to non-satellite sequence insertions compared to centromeric regions. Furthermore, while inversions are present in centromeric and pericentric regions, they are more frequent within centromeric regions. The higher frequency of inversions in centromeric regions is likely due to their more homogeneous sequence composition than the more variable pericentric regions. Our study shows the progressive increase in non-satellite repeat density from centromeres to pericentromeres to chromosomal arm junctions. These non-satellite repeats, including retrotransposons, LINEs, and SINEs, play crucial roles in genome integrity and genetic diversity. This gradient of repeat density may reflect the complex structural organization required for the transition between different chromosomal domains.

We found that core centromeric regions occupied by 120-mer MiSats exhibit high levels of CENP-A, a crucial protein for centromere function. However, MiSat length variants, including 112-mers and 112-64-dimers, located at the centromere and pericentromeric junctions of a few centromeres, displayed reduced CENP-A levels. Interestingly, these variants contain a higher density of CENP-B boxes than the 120-mer MiSats, and we observed corresponding increased CENP-B enrichment on these minor satellite length variants despite their lower levels of CENP-A. In addition to interacting with CENP-A, CENP-B has been shown to associate with heterochromatic proteins such as HP1 and Suv39h1 (Otake et al., 2020). Our observation that CENP-B is enriched at centromeric-pericentric transition regions suggests that its interaction with heterochromatin proteins is influenced by its proximity to pericentric regions. However, why this pattern is specific to a few chromosomes remains unclear. We also found that the chromatin assembled at the pericentric regions varies depending on the type and location of MaSats. Homogeneous MaSats, predominantly localized near centromeres and in the core regions of the pericentromeres, were more enriched with H3K9me3, a marker of transcriptionally silent heterochromatin. In contrast, divergent MaSats, concentrated at the pericentromeric-chromosomal arm transitions, exhibited lower levels of H3K9me3. Interestingly, we observed the opposite pattern of H3K27me3 enrichment on MaSats. Homogeneous MaSats showed lower enrichment of H3K27me3, while divergent MaSats and non-satellite islands within MaSat regions were more enriched with H3K27me3. In humans, the centromeric core shows a sharp dip in DNA methylation (Altemose et al., 2022). While we observed a decrease in DNA methylation within mouse centromeric regions compared to the pericentric regions, this reduction was not as pronounced as in humans. In contrast, the pericentric regions in mice displayed highly dense levels of DNA methylation. Additionally, we found that only a small subset of MaSat and MiSat sequences were transcribed into RNA. Within homogeneous MaSats, RNA expression was associated with regions of lower DNA methylation density, suggesting a possible link between reduced methylation and transcriptional activity.

In conclusion, our study provides comprehensive genomic and epigenomic maps of mouse centromeric, pericentromeric, and adjacent junction regions using PacBio long-read sequencing (summarized in Figure 7). This work lays the foundation for future research aimed at producing a complete telomere-to-telomere (T2T) assembly of the mouse genome, similar to the recent T2T assembly of the human genome.

**Figure 7.**
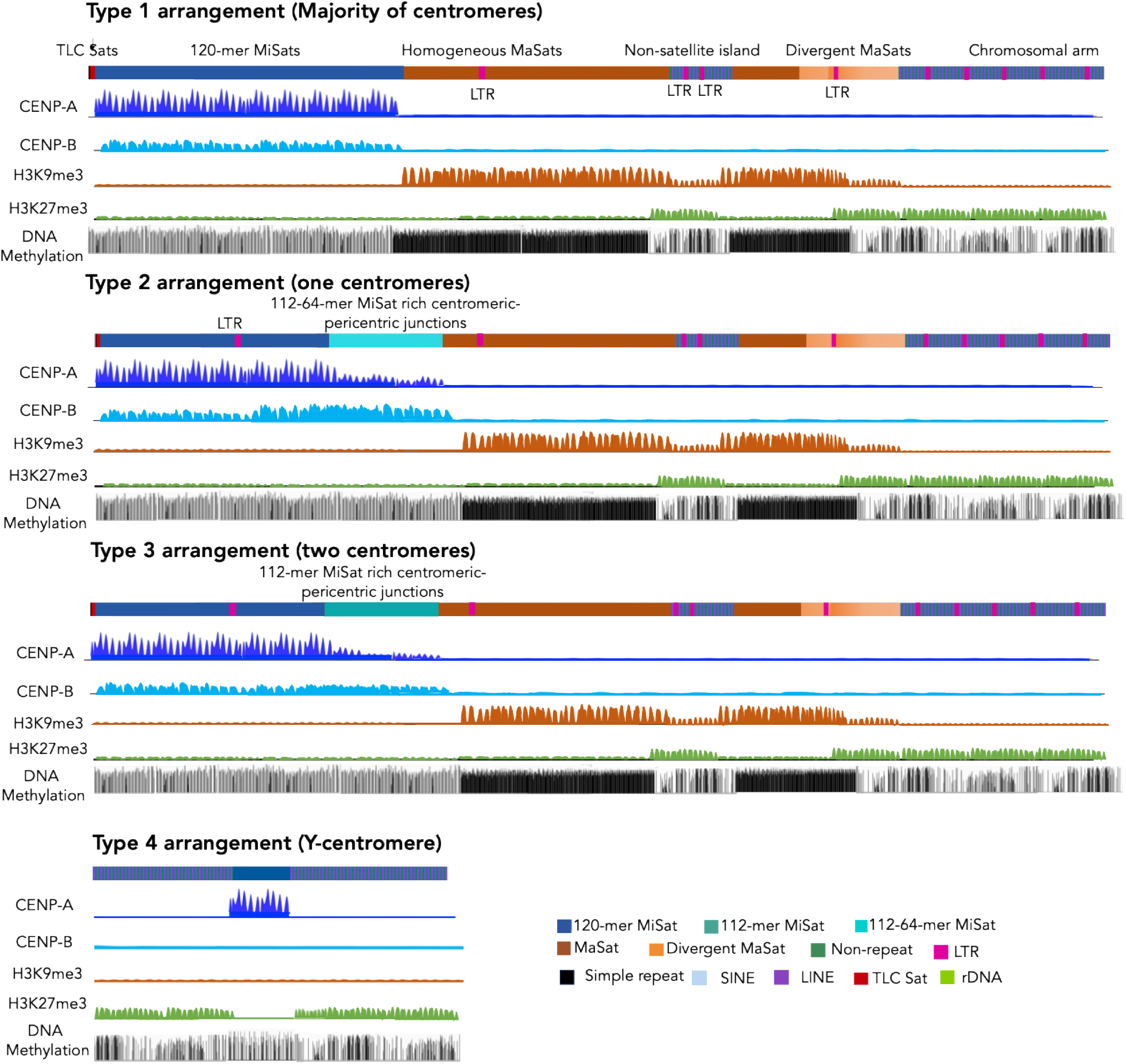
Representative maps of genetic and epigenomic features of mouse centromere and pericentromeres.

## Material and methods

### Animals

C57BL/6J animals were purchased from Jackson Laboratory and maintained at Emory University per the protocol approved by Institutional IACUC.

### Nuclei preparation

Nuclei were prepared from C57BL/6J kidney tissues. First, the frozen tissue was weighed and ground into a fine powder using liquid nitrogen in a prechilled mortar and pestle. The powdered tissue was then resuspended in 4 mL of ice-cold extraction Buffer (0.32 M sucrose, 60 mM KCl, 15 mM NaCl, 15 mM Tris-Cl (pH 7.5), 5 mM MgCl2, 0.1 mM EGTA, 0.5 mM DTT, and 1X protease inhibitor) per gram of tissue. The suspension was homogenized using a Tekmar tissuemizer for 15 pulses of approximately 5 seconds each, with 20-second intervals on ice. Finally, the cell suspension was filtered through a 100 μm cell strainer to remove large tissue chunks. The filtered homogenate suspension was centrifuged at 6000 x g for 10 minutes at 4°C. The pellet was resuspended in 4 mL (per gram of tissue) of ice-cold Extraction buffer supplemented with 0.2% IGEPAL, and the samples were incubated on ice for 10 minutes. The nuclei suspension was layered on top of ice-cold sucrose buffer (1.2 M sucrose, 60 mM KCl, 15 mM NaCl, 15 mM Tris-Cl (pH 7.5), 5 mM MgCl2, 0.1 mM EGTA, 0.5 mM DTT, and 1X protease inhibitor) to create a sucrose cushion. Using prechilled rotors, the mixture was centrifuged at 10,000 x g for 20 minutes at 4°C. The nuclei pellet was resuspended in a buffer suitable for downstream processing.

### PacBio sequencing

High molecular weight genomic DNA was isolated from the kidneys of C57BL/6J mice using the Monarch High Molecular Weight DNA Extraction Kit, following the manufacturer’s instructions. The nuclei pellet was incubated with 600 ul Monarch Lysis Buffer and 20 ul Proteinase K and incubated at 55°C with gentle agitation at 700 rpm for 45 minutes. Following lysis, 10 μL of RNase A was added to the lysate and incubated at 55°C with gentle agitation at 700 rpm for 45 minutes for 10 minutes. Subsequently, 300 ul of Monarch Protein separation Buffer was added to the lysate, mixed vigorously for 1 minute by inverting the tube and centrifuged at 16,000 x g for 10 minutes to pellet the precipitated proteins.

The DNA supernatant was transferred to a clean tube containing 500 ul isopropanol and two Monarch DNA capture beads. The solution was mixed by vertically rotating the tube at 10 rpm for 5 minutes to attach DNA to the beads. The supernatant was discarded, beads were washed twice with 500 μl Monarch gDNA wash buffer, and DNA was eluted from beads with 200 μL of Monarch DNA Elution Buffer. The purified high molecular weight genomic DNA was sequenced using PacBio's Revio platform with single-molecule, real-time (SMRT) sequencing technology to generate long reads.

Raw sequence data generated by the Revio platform was processed using PacBio's SMRT Analysis software suite. This included base calling, quality filtering, and assembly of high-fidelity (HiFi) reads. The HiFi reads, known for their high accuracy, were used for downstream genomic analyses, including genome assembly, structural variant detection, and methylation analysis. This comprehensive approach ensured the generation of long, high-quality reads suitable for in-depth genomic studies.

### CUT&RUN sequencing

The pelleted nuclei were resuspended in CUT&RUN wash buffer (20 mM HEPES (pH 7.5), 150 mM NaCl, and 0.5 mM spermidine). Concanavalin beads were prepared by washing the bead slurry with 3 times the volume of binding buffer (20 mM HEPES, 10 mM KCl, 1 mM CaCl2, and 1 mM MnCl2) and resuspended in an equal volume of binding buffer as the initial slurry volume.To each CUT&RUN reaction, 20 μL of beads per million nuclei were added and the mixture was incubated at room temperature for 15 minutes on a rotator. The beads were washed, resuspended in 200 μL of antibody buffer containing 4 μL of antibodies per reaction, and incubated at 4°C overnight. The beads were washed in 500 μL of cold Dig-Wash buffer (20 mM HEPES (pH 7.5), 150 mM NaCl, 0.5 mM spermidine, and 0.05% digitonin) twice, resuspended in 200 μL of Dig-Wash buffer containing 1ul pA-MN, and incubated for 1 hour at room temperature on a rotator. The beads were then washed twice with cold Dig-Wash buffer, resuspended in 200 μL of Dig-Wash buffer with 3mM CaCl2, and incubated at 0°C for 30 minutes. After 30 minutes, the reaction was stopped by adding 200 μL of 2X DRSTOP buffer (550 mM NaCl, 20 mM EDTA, 4 mM EGTA, 0.05% digitonin, 40 μg/mL glycogen, 50 mg/mL RNase A and 0.0025 ng/μL yeast spike-in DNA) and the mixture was incubated for 10 minutes at 37°C. The samples were centrifuged for 5 minutes at room temperature, and the supernatant was collected. DNA was extracted from the supernatant and libraries were prepared using KAPA HyperPrep kit. Amplified libraries were sequenced using NextSeq 500/550 instrument.

### Generation and Annotation of Satellite Contigs

Assembled contigs were generated from PacBio HiFi reads using Hifiasm software with default parameters. Satellite-containing contigs were then filtered from the total contig pool using NCBI-BLAST against satellite consensus sequences. Specific satellite types, including 120-mer MiSats, 112-mer MiSats, 112-64-mer MiSats, TLC-sats, homogeneous MaSats, and divergent MaSats, were annotated on these satellite contigs using NCBI-BLAST. Non-satellite repeat sequences (LINE, SINE, LTR, simple repeats, rDNA, and tRNA) were annotated using RepeatMasker on *Mus musculus* repeat database (Smit, 2013-2015).

### Annotations of Genes in Pericentric-Chromosome Arm junctions

We identified 2-5 kb sequences from the extreme pericentric end of each chromosome as annotated in the GRCm39 genome. These sequences were subsequently used in NCBI-BLAST searches against Hifiasm contigs to identify contigs containing the respective sequences. Next, genes and transcribed pseudogenes located within the first 5Mb (if the contig is >5Mb) of all autosomes except chromosomes 2, 3, and 10, as well as Chromosome X scaffold NT_165789.3, were identified using the NCBI genome annotation associated with the mouse GRCm39 genome. Genes specific to Chromosome Y near the centromere were similarly identified.

The corresponding RNA transcripts were searched for matches against the Hifiasm contigs using NCBI BLAST. BLAST hits with greater than 99% identity were selected for further analysis. For each chromosome, the selected hits were further restricted to the previously identified contig containing the pericentric junction. This contig was converted to a BED file and loaded into IGV for visual comparison to GRCm39 (viewed using the NCBI Genome Data Viewer). Non-specific BLAST hits were identified by comparison to GRCm39 and removed. The resulting cleaned BED file was then processed to combine all exon BED records into a single BED record per gene.

### DNA methylation analysis

To analyze DNA methylation, Bam files from PacBio HiFi reads containing DNA methylation kinetics were mapped to satellite-containing Hifiasm contigs using the pbmm2 aligner, which is part of the CCSmeth pipeline, using default settings. The resulting mapped Bam files were then processed to extract methylation information using pb-CpG-tools, which was run in the “count” pileup mode setting to generate CpG scores. The CpG scores were then converted into BigWig files for visualization. The generated BigWig files were loaded into the Integrative Genomics Viewer (IGV) for visual analysis.

### RNA-sequencing and analysis

RNA was isolated from C57BL/6J mouse kidney and sequenced by Azenta Life Science. The quality of sequenced reads was assessed using FastQC. Poor-quality regions and adapter sequences were trimmed using Trimmomatic. Resulting sequencing reads were directly mapped to the Hifiasm satellite contigs using STAR(v2.7.3a) (Dobin et al., 2013) aligner with default settings. The aligned bam files were used to calculate expression levels as transcripts per million and were also visualized by IGV (v2.16.1).

## Acknowledgment

We thank Thakur lab members for their discussions and input on the manuscript. Illumina sequencing of CUT&RUN samples was performed at Emory Integrated Genomics Core. PacBio sequencing and RNA sequencing were performed at Azenta Inc.

## Notes

### Competing Interest Statement

The authors have declared no competing interest.

