## Supplementary Figures for "Genomic and epigenomic maps of mouse centromeres and pericentromeres"

### Supplementary Information

### Supplementary Figure 1

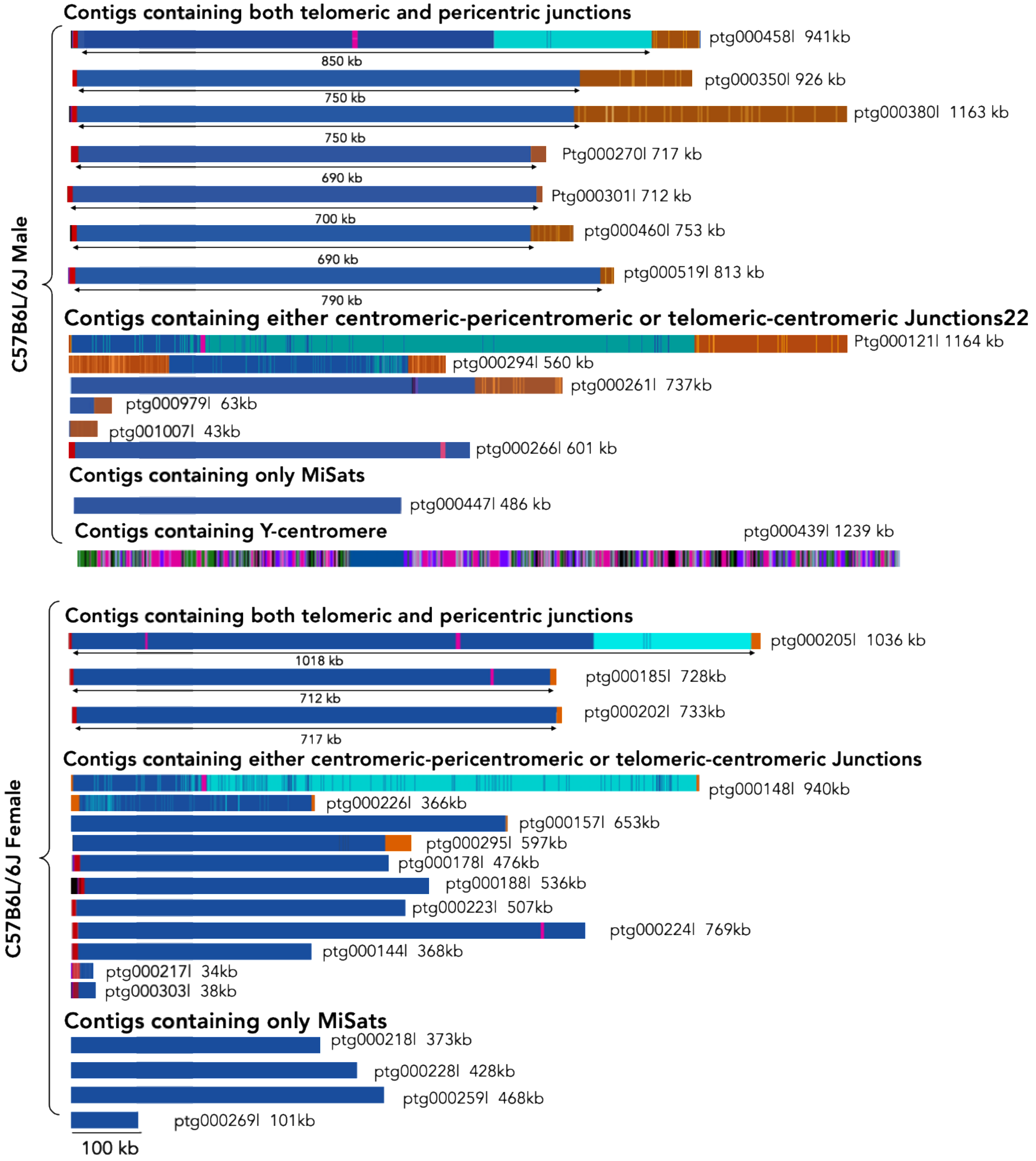


**Supplementary Figure 1:** All MiSat containing contigs in male (this study, Top) and female (Hon et al., 2020) (Bottom) C57BL/6J mice.

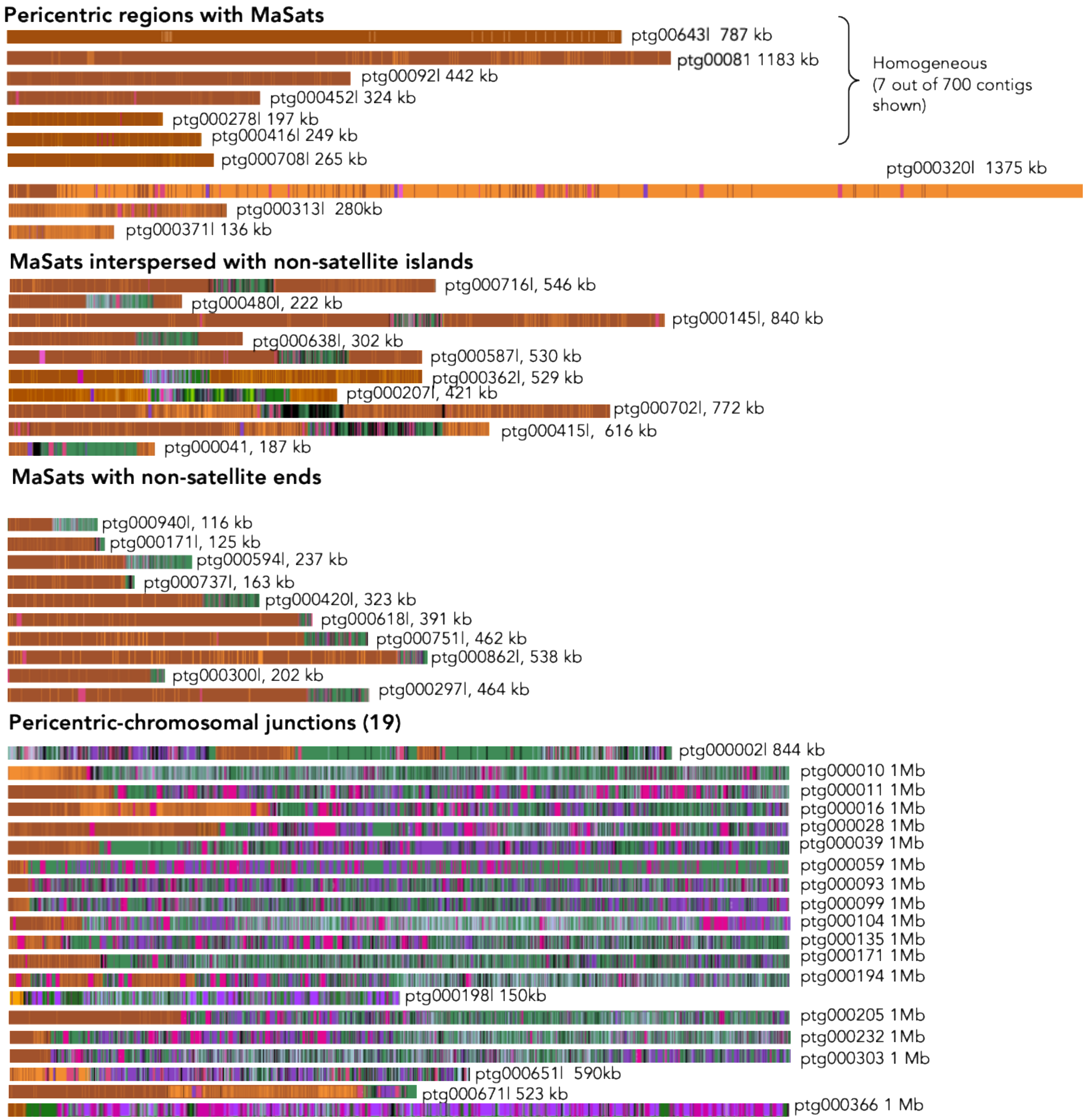


**Supplementary Figure 2:** Assemblies for pericentric and pericentric-chromosomal arm junctions.

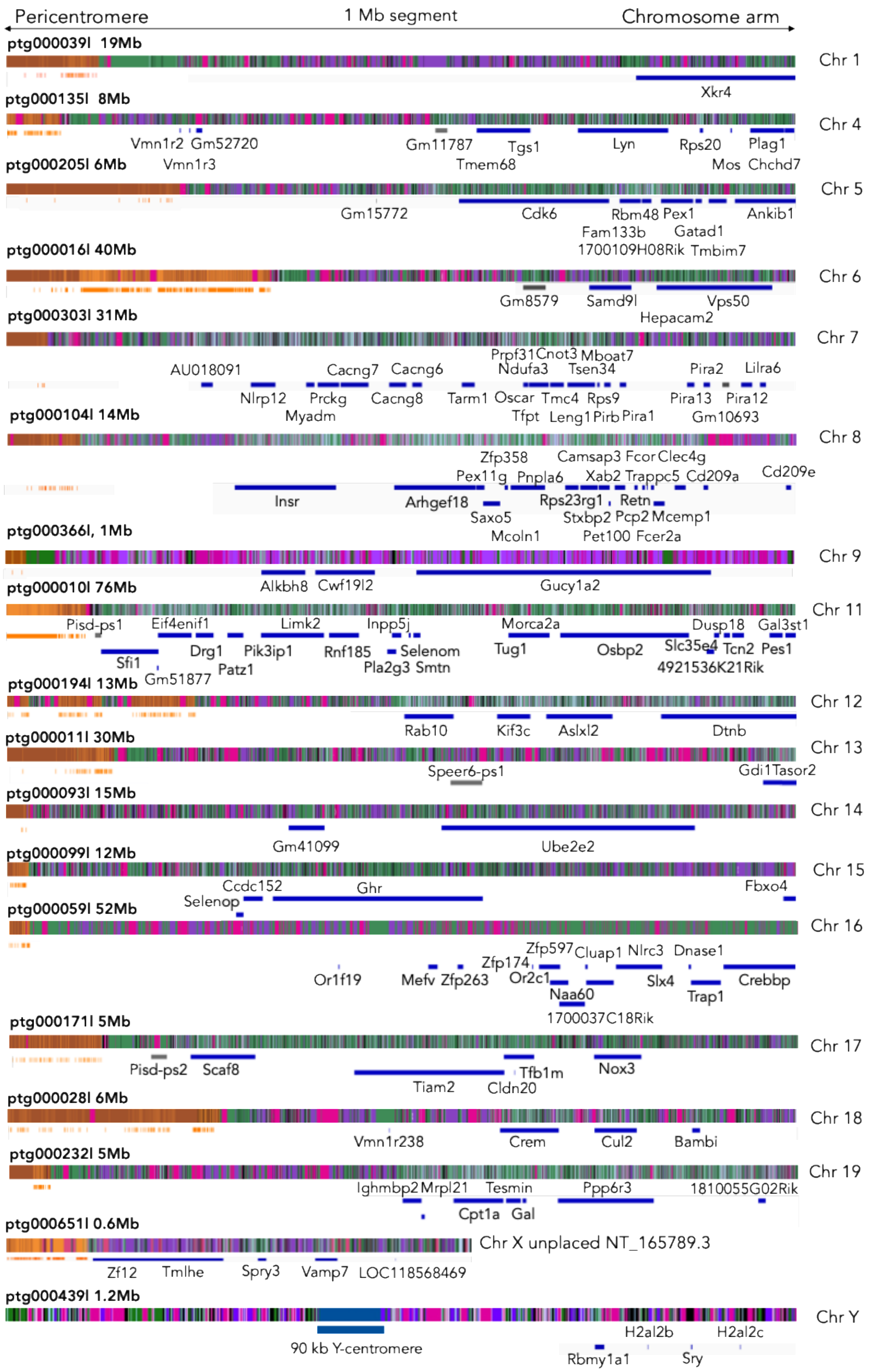


**Supplementary Figure 3:** Assemblies maps of pericentric-chromosomal arm junctions. Orange, blue, and grey bars below each contig represent divergent MaSats, genes and pseudogenes, respectively. All chromosome-specific genes and pseudogenes we identified in these junctions are listed.

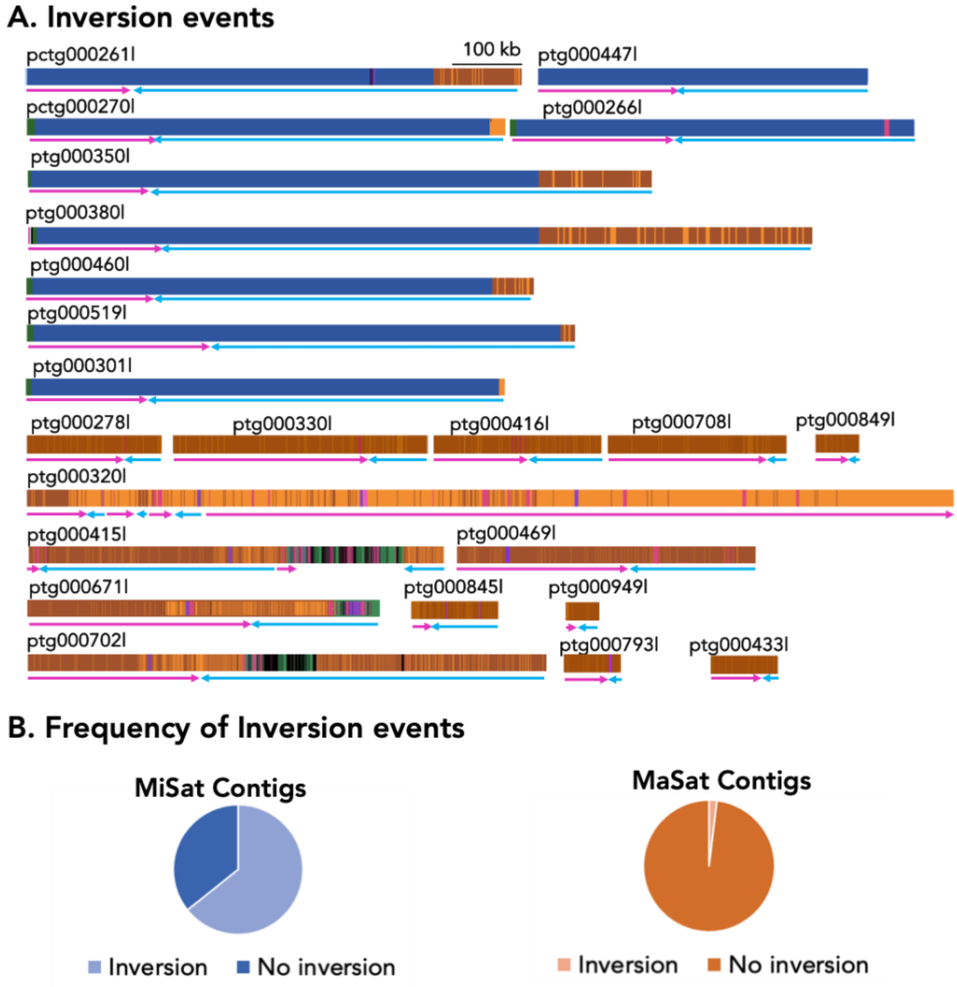


**Supplementary Figure 4. Inversion events and the sequence motif density at centromeric and pericentromeric regions. A.** All inversion events in centromeric and pericentromeric contigs are shown. A change in the direction and color of the arrow below a given contig indicates an inversion. **B.** Frequency of inversion events in MaSat and MiSat contigs.

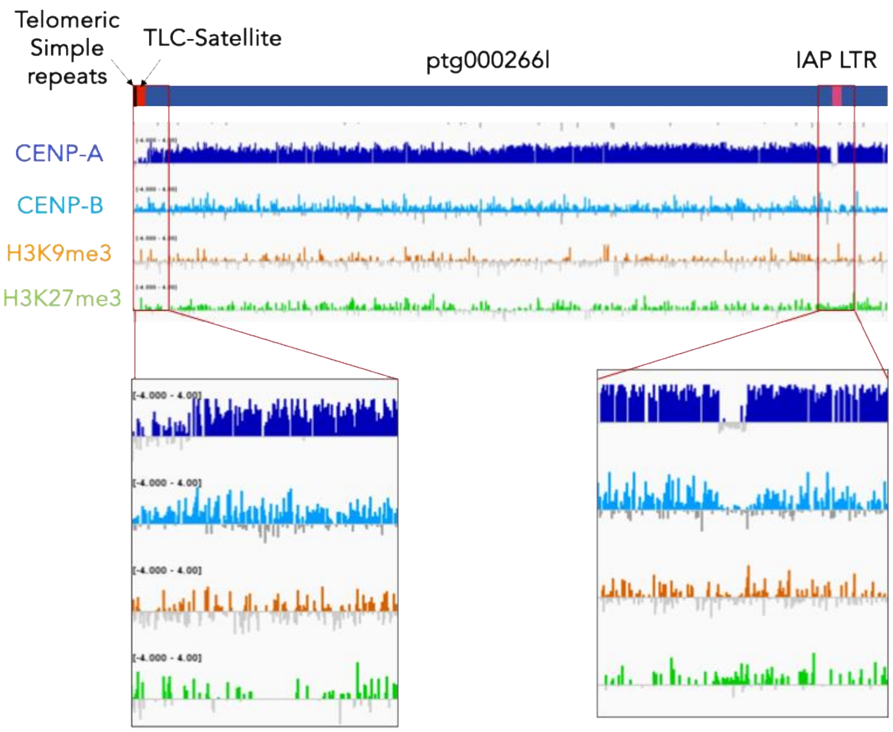


**Supplementary Figure 5.** Chromatin profiles on centromere-telomeric junctions and interspersed transposons within MiSat contigs
